# A defined medium based on R2A for cultivation and exometabolite profiling of soil bacteria

**DOI:** 10.1101/2021.05.23.445362

**Authors:** Markus de Raad, Yifan Li, Peter Andeer, Suzanne M. Kosina, Nicholas R. Saichek, Amber Golini, La Zhen Han, Ying Wang, Benjamin P. Bowen, Romy Chakraborty, Trent R. Northen

**Author notes:** Corresponding Author Trent R. Northen, Lawrence Berkeley National Laboratory, 1 Cyclotron Road, Berkeley, CA 94720, USA.

## Abstract

Exometabolomics is an approach to assess how microorganisms alter their environments through the depletion and secretion of chemical compounds. Comparisons of inoculated with uninoculated media can be used to provide direct biochemical observations on depleted and secreted metabolites which can be used to predict resource competition, cross-feeding and secondary metabolite production in microbial isolates and communities. This approach is most powerful when used with defined media that enable tracking of all depleted metabolites. However, microbial growth media have traditionally been developed for the isolation and growth of microorganisms but not metabolite utilization profiling through LC-MS/MS. Here, we describe the construction of a defined medium, the Northen Lab Defined Medium (NLDM), that not only supports the growth of diverse bacteria but is defined and therefore suited for exometabolomic experiments. Metabolites included in NLDM were selected based on their presence in R2A medium and soil, elemental stoichiometry requirements, as well as knowledge of metabolite usage by different bacteria. We found that NLDM supported the growth of 53 phylogenetically diverse soil bacterial isolates and all of its metabolites were trackable through LC–MS/MS analysis. These results demonstrate the viability and utility of the constructed NLDM medium for cultivating and characterizing diverse microbial isolates and communities.

**Originality-Significance Statement:** We build a defined medium based on the metabolite composition of R2A medium and soil, elemental stoichiometry requirements, and knowledge of metabolite usage by different bacteria. The newly formulated defined medium was evaluated on its ability to support the growth of soil isolates and its application for metabolite utilization profiling. We found that of 53 phylogenetically diverse soil bacterial isolates grew on the defined medium and all of its metabolites were trackable through LC–MS/MS analysis. This demonstrates the viability and utility of the constructed defined medium for cultivating and characterizing diverse microbial isolates and communities.

## Introduction

Exometabolomics, or metabolic footprinting, is an approach to determine the metabolites produced or depleted in a given environment(Allen *et al*., 2003). For example, an exometabolomic experiment, may use Liquid Chromatography Tandem Mass Spectrometry (LC-MS/MS) to compare growth media before and after microbial growth to identify metabolites that a given microorganism, or microbial community, secrete and consume under specific growth conditions(Kosmides *et al*., 2013).

While any growth media can be used for exometabolite profiling, defined media are desirable as their composition is more trackable and tractable. In addition, complex media components are frequently derived from complex organisms (e.g. yeast extract) that can vary in relative composition between batches. Thus, defined media enable a complete view of trade-offs in substrate use and allow differentiation between secreted products and the breakdown of complex substrates by extracellular enzyme activities on biopolymers. For example, sugars resulting from extracellular polymer degradation could be misinterpreted as secreted products.

Many different media exist for culturing soil bacteria, for example the selective culturing of environmental microbes. These culture media can be grouped as rich media or defined media. For instance, Reasoner’s 2A (R2A) medium is one of the most widely used nutrient-rich media for isolating and culturing soil microbes and defined media designed for isolation often have a single substrate or nutrient source to select for specific organisms(Reasoner and Geldreich, 1985; Vartoukian *et al*., 2010; Trinh *et al*., 2019). Although R2A medium was not designed to be ecologically relevant for soil environments, it has been found to support the growth of a wide variety of soil microbes (Reasoner and Geldreich, 1985; Marteinsson *et al*., 2015; Nguyen *et al*., 2018; Chaudhary *et al*., 2019). Previously, we reported the development of a defined medium based on water soluble soil metabolites from saprolite soil (Jenkins *et al*., 2017). Although it was successfully used for metabolomic profiling, it was found to support the growth of only half as many isolates as R2A medium.

Here, we describe the construction of another defined medium, Northen Lab Defined Medium (NLDM), designed for exometabolomic analysis of soil bacteria while supporting the growth of diverse bacteria. All metabolites and their relative abundances included in NLDM were selected on the basis of 1) their presence in R2A medium, soil and other environments, 2) knowledge of elemental stoichiometries for bacterial growth(Cleveland and Liptzin, 2007) and 3) existing exometabolomic data on substrate use across diverse bacteria(Reasoner and Geldreich, 1985). To validate NLDM as both a viable growth substrate and a valuable exometabolomic medium, a panel of 53 phylogenetically diverse isolates from the Oak Ridge Field Research Center (ORFRC) were grown in NLDM and R2A to compare their ability to support bacterial growth and 7 isolates were analzyed on substrate usage.

## Results

NLDM is primarily composed of Wolfe’s minerals, Wolfe’s vitamins, and the metabolite composition of R2A, to balance microbial growth, metabolite diversity, and compositional simplicity (avoiding an excessive number of metabolites). In addition, the exometabolomic assertion repository, Web of Microbes, was used to assess if the included metabolites are commonly consumed by bacteria(Kosina *et al*., 2018). Metabolite concentrations in NLDM were adjusted to 1) have the C:N ratio consistent with the soil microbial biomass and 2) mimic compound class ratios found in R2A(Cleveland and Liptzin, 2007).

### Metabolite selection

Except for glucose and pyruvic acid, R2A is a complex and undefined metabolite mixture, primarily based on yeast extract and proteose peptone(Reasoner and Geldreich, 1985). To identify the small molecules from these complex components, we previously analyzed R2A medium and several different soils using LC-MS/MS and gas chromatography MS (Supplementary Table 1)(Liebeke *et al*., 2009; Swenson *et al*., 2015; Jenkins *et al*., 2017; Kosina *et al*., 2018; Sasse *et al*., 2019). Through these efforts, a number of metabolites present in R2A and soils were identified. These included most of the standard amino acids, all 5 standard nucleobases, and 3 standard ribonucleosides. Based on these findings, we included all 20 standard amino acids, the 5 standard nucleobases, and the 5 standard ribonucleosides in NLDM. In addition, the nucleobases and ribonucleosides xanthine, hypoxanthine, inosine and xanthosine were included in NLDM based on their presence in R2A and soils.

Primary energy sources in R2A are the sugar glucose and the glucose polymer starch; the latter is too large for small molecule LC-MS/MS detection. As a result, we included glucose plus two additional sugars detected in R2A: the dihexose trehalose and the sugar alcohol myo-inositol. In addition, to increase substrate diversity and assess additional metabolic pathways, we also included the pentose xylose, and the amino sugar N-acetyl-glucosamine, which are commonly found in soils(Gunina and Kuzyakov, 2015; Ni *et al*., 2020).

Pyruvic acid is another major defined energy source in R2A medium. To capture organic acids as potential energy sources for bacteria pyruvic acid was included along with 6 other common organic acids detected in R2A medium and/or in soil (Supplementary Table 1). Seventeen other metabolites were selected for NLDM based on analysis of R2A medium and/or soil.

We decided to include spermidine even though it was not detected in R2A or soil because polyamines are ‘essential’ cofactors and because we had detected another polyamine, A-acetylputrescine, in R2A medium(Xavier *et al*., 2017). Also, the amino sugar n-acetylmuramic acid was not detected in R2A or the soil samples, but we included it since it had been previously found in soils and is a component of bacterial cell walls(Glaser *et al*., 2004; Ni *et al*., 2020). Vitamins were not included in the formulation of NLDM as they will be added separately. In total, 64 metabolites were selected to be included in NLDM (Supplementary Table 2).

### Mining existing exometabolite data for refining NLDM formulation

After the formulation of NLDM, we checked if the selected metabolites can be used by microbes. To do this, we analyzed the usage of the selected metabolites by microbes using existing exometabolomic data collected in Web of Microbes(Kosina *et al*., 2018). A metabolite was deemed used/converted if the metabolite was significantly lower in the presence of a microbe compared to the uninoculated control. Out of the 64 metabolites present in NLDM, 54 were in the Web of Microbes database. All but 4 metabolites, α-ketoglutaric acid, cysteine, cytidine and uridine, were used/converted by at least 1 microbe (Supplementary Table 3).

### NLDM formulation

The quantitative formulation of the 64 selected metabolites was based on the amount of organic carbon (C) and nitrogen (N) in R2A, calculated as 1146 mg/L and 144 mg/L, respectively(Kim *et al*., 2019). We divided all 64 metabolites in NLDM into 4 different groups: sugars, organic acids, amino acids and other metabolites (Supplementary Table 2). Metabolites within each group were assigned fixed equimolar concentrations and the total organic C and N was calculated (Table 1). This yielded a C:N ratio of 9:1, which is similar to R2A (10:1) and the soil microbial biomass ratio (9:1)(Cleveland and Liptzin, 2007). As for the salts, NLDM contains 5 mM phosphate, 1 mM ammonium, 2 mM sodium, 7 mM potassium, 1 mM magnesium, 1 mM sulfur, 1 mM calcium and 2 mM chloride (Supplementary Table 2). NLDM is supplemented with 1x Wolfe’s vitamins and 1x Wolfe’s minerals.

**Table 1.** Breakdown of organic C and N in NLDM.

|  | uM of each metabolite | # of compounds | C (mg/L) | N (mg/L) |
| --- | --- | --- | --- | --- |
| Sugars | 875 | 7 | 578 | 12 |
| Organic Acids | 525 | 7 | 195 | 22 |
| Amino Acids | 175 | 20 | 225 | 71 |
| All other metabolites | 17.5 | 30 | 43 | 18 |
|  | **Total** | **64** | **1042** | **123** |

### NLDM is a viable medium for exometabolite profiling

To confirm that NLDM is suitable for exometabolomic studies, the substrate preferences of 7 different bacterial isolates were determined using LC-MS/MS. The isolates included 4 *Pseudomonas* species (FW305-3-2-15-E-TSA4, GW531-R1, GW456-L15 and FW305-3-2-15-C-LB3), the *Cupriavidus* sp. FW305-77, the *Delftia* sp. GW456-R20 and the *Sphingopyxis* sp. GW247-27LB. To comapre substrate preference between isolates from the same genus, 4 *Pseudomonas* isolates were selected. The other isolates were selected based on their growth profiles, which were different to the *Pseudomonas* isolates growth profiles. Using HILIC LC-MS/MS, all metabolites were detected in the full medium formulation (Supplementary Table 4). Clear differences in metabolite depletion were observed both between isolates and between metabolite classes (Figure 1). All 64 metabolites from the NLDM medium were utilized by at least 1 isolate after 24 h (compared to medium control using ANOVA with Tukey’s honestly significant difference test, *P* < 0.05). *Pseudomonas* GW531-R1 and FW305-3-2-15-E-TSA4 used the most metabolites (57 significantly depleted) whereas *Cupravidius* FW305-77 used the fewest (45). Hierarchical clustering revealed multiple distinct clusters. Cluster 2 consists of the most depleted compounds, and contains most of the nucleobases and nucleosides. Clusters 1 and 3 contain the least depleted compounds. Clusters 4 and 5 are composed of intermediate depleted metabolites and contain most of the amino acids and organic acids. All 4 *Pseudomonas* species had similar utilization compared to the other isolates.

**Figure 1.**
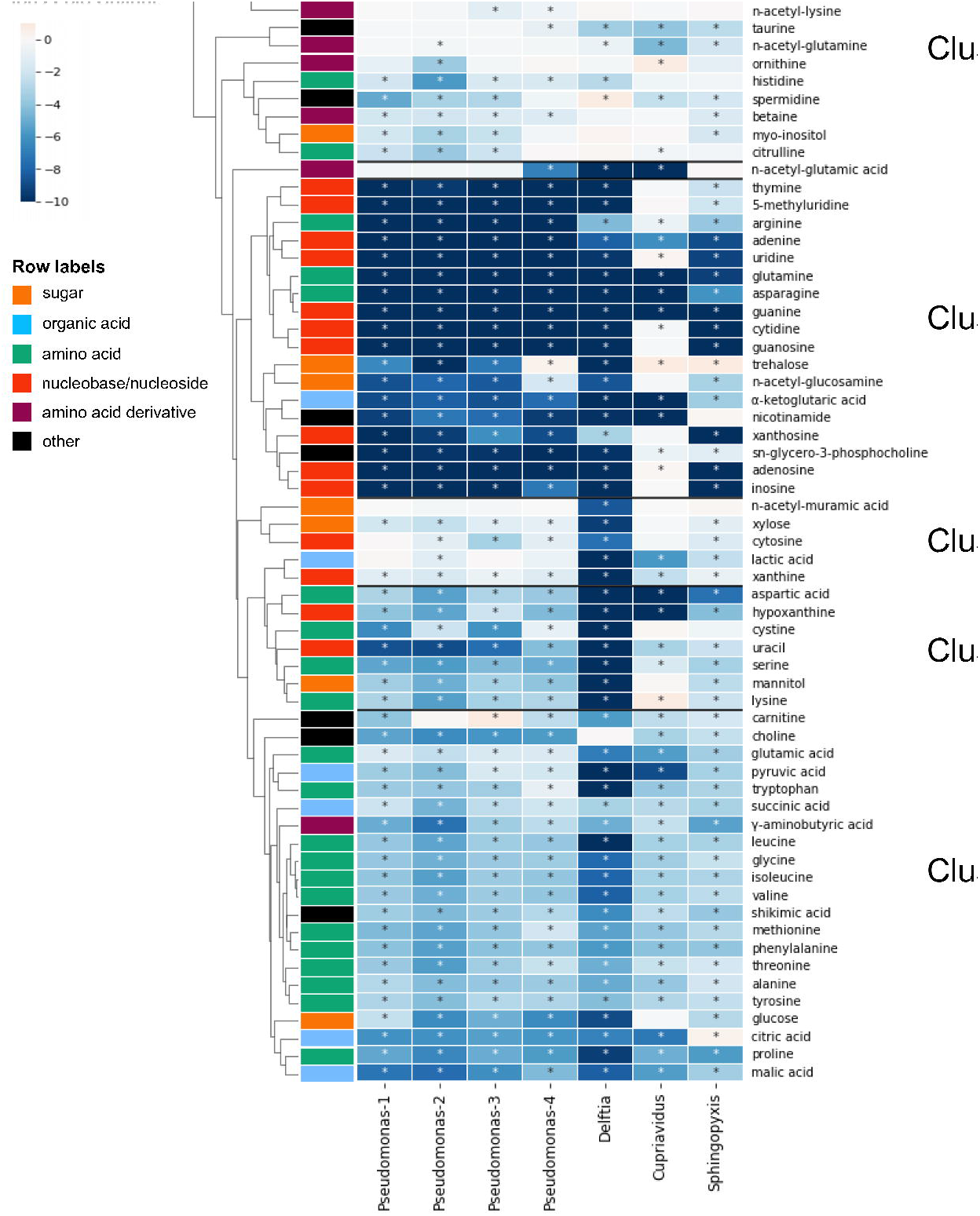
NLDM metabolite utilization patterns by soil isolates. NLDM metabolites utilization after 24 hours displayed as the log2 of the average normalized peak height or peak area relative to the medium control in a clustering heatmap. n□=□4 for each isolate. Metabolite row groups are colored according to their chemical class (orange = sugar, sky blue = organic acid, bluish green = amino acid, vermillion = nucleoside/nucleobase, reddish. Strains are abbreviated as follows: ‘Pseudomonas-1’ *Pseudomonas* sp. FW305-3-2-15-C-LB3, ‘Pseudomonas-2’ *Pseudomonas* sp. GW531-R1, ‘Pseudomonas-3’ *Pseudomonas* sp.GW456-L15, ‘Pseudomonas-4’ *Pseudomonas* sp.FW305-3-2-15-E-TSA4, ‘Delftia’ *Delftia* sp. GW456-R20’, ‘Cupriavidus’ *Cuprividus* sp. FW305-77, ‘Sphingopyxis’ *Sphingopyxis* sp. GW247-27LB’. * indicates *P* < 0.05 (ANOVA with Tukey’s honestly significant difference test).

**Figure 2.**
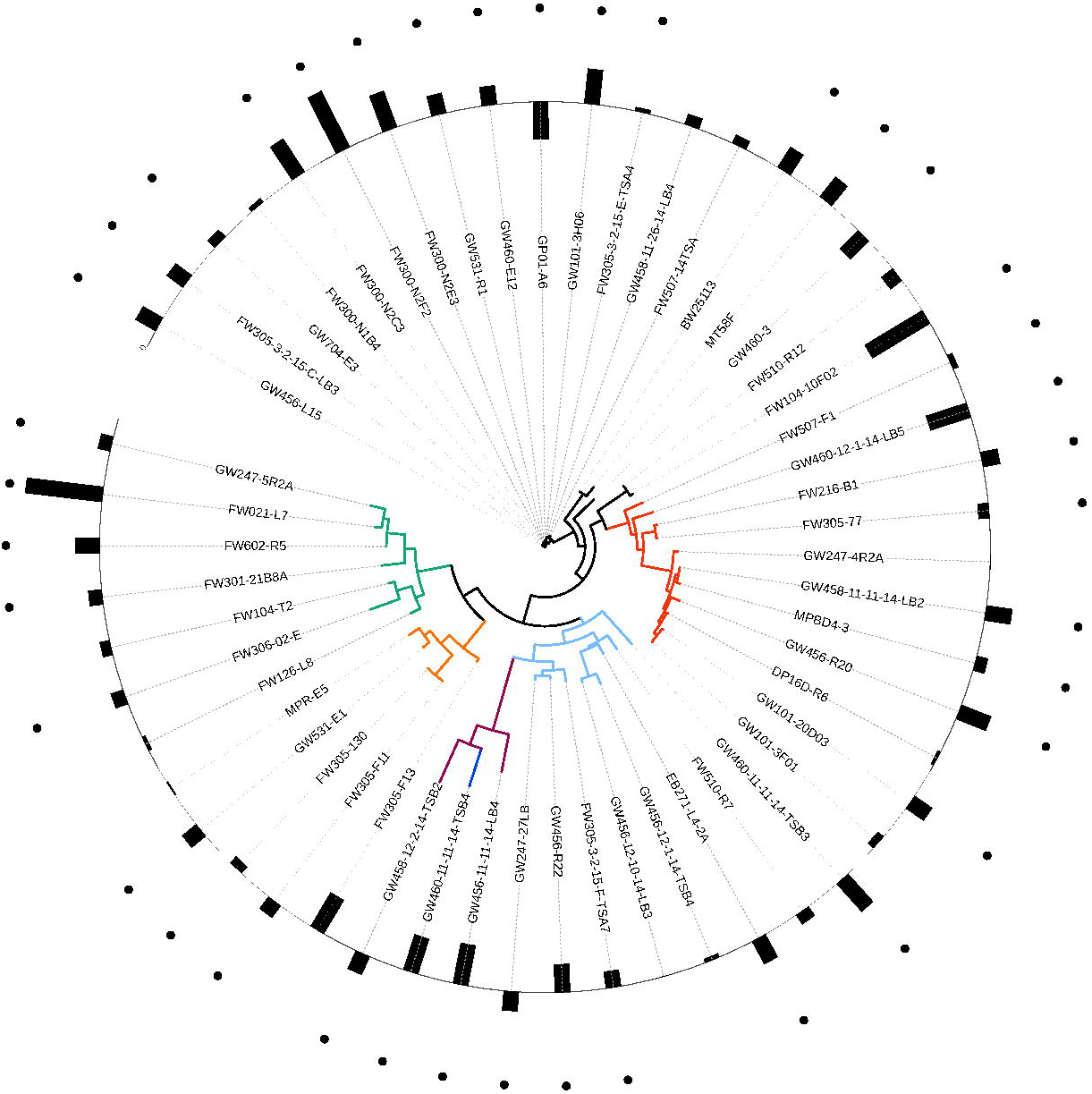
Phylogenetic tree of all isolates with corresponding ratio of biomass yield on NLDM and R2A. Branch colors indicate the phylogenetic origin of each isolate by class (orange = Actinobacteria, sky blue = Alphaproteobacteria, bluish green = Bacilli, vermillion = Betaproteobacteria, reddish purple = Flavobacteria, black = Gammaproteobacteria, blue = Sphingobacteriaceae). Bars on the outer circle indicate the average (n=3) log2 ratio of the biomass yield (maximum OD_600_) of each isolate grown on NLDM and R2A. Ratios >0 indicate that biomass yield on NLDM was larger than on R2A and ratios <0 indicate that that biomass yield on R2A was larger than on NLDM. Black dots indicate significant difference (two-sample *t*-test, *P* < 0.01). All growth data (OD_600_ values over time for all isolates) can be found in Supplementary Table 5A and 5B.

### Comparable growth is observed on NLDM and R2A

To determine if NLDM was suitable for the growth of diverse soil bacteria, 53 bacterial isolates from the ORFRC were grown in both NLDM and R2A (R2A was used to isolate these isolates). It was found that all of the isolates grew in both media (Supplementary Table 5A and 5B). The overall biomass yield (as measured as the highest observed OD_600_ after background correction) was similar for R2A and NLDM (Supplementary Figure 1). While individual isolates exhibited significant differences in biomass yields on one medium vs the other, no significant differences between the biomass yields across phylogenetic classes or families were observed for NLDM vs R2A. Isolate growth rates were also similar on the two media (Supplementary Figure 1 and 2). There were no significant differences between the growth rates of different phylogenetic classes or families on NLDM as compared with R2A.

## Discussion

The goal of this study was to develop a defined medium suitable for both the cultivation and exometabolite profiling of diverse soil bacteria. R2A, which was developed for the isolation and growth of oligotrophic, environmental bacteria and is perhaps the most widely used medium for that purpose(Nishioka *et al*., 2016). For this reason, the composition of R2A was used to guide the NLDM formulation. We additionally considered metabolites previously detected in soil, existing isolate exometabolomics data, elemental stoichiometry in refining the media formulation, and limited the number of metabolites to balance relevance with the cost and time required to prepare the media. NLDM supported the growth of all screened bacterial isolates and all the metabolites included in NLDM were traceable by LC-MS/MS.

Most of the metabolites included in NLDM are also present in several other soil-extract based media(Liebeke *et al*., 2009; Jenkins *et al*., 2017; Nguyen *et al*., 2018; Sasse *et al*., 2019). A major difference between these media and NLDM is that NLDM contains a single phospholipid whereas other soil-extract media contain a range of different fatty acids. The metabolite diversity in NLDM is lower than rich media, which include undefined and variable components such as yeast extract. However, we determined this is a worthwhile tradeoff given that this media allows for a comprehensive view of metabolite use using LC-MS/MS. We anticipate that NLDM can be used as a base medium for supplementation with other compounds of interest to increase its relevance to soil and to extend to additional classes of metabolites.

The final concentrations of sugars, organic acids, amino acids, and other metabolites included in NLDM make up a C:N ratio that is consistent with R2A and the “redfield ratio” for soil bacterial biomass(Cleveland and Liptzin, 2007). Compared to soil bacterial biomass, the phosphate concentration in NLDM is high to ensure that phosphate is not limited during culturing. We acknowledge that the chosen metabolite concentrations do not reflect ecological soil conditions, as metabolite concentrations in soil vary by type. Like R2A, NLDM is a nutrient rich medium. This can be an issue since it is thought that nutrient rich culture media favor the growth of faster-growing bacteria at the expense of slow-growing species which can also be inhibited by substrate-rich conventional media(Vartoukian *et al*., 2010; Nunes da Rocha *et al*., 2015; Overmann *et al*., 2017; Bartelme *et al*., 2020). We anticipate that NLDM can also be diluted to examine more oligotrophic bacteria, however, the culture volume will need to be increased to provide sufficient signal for LC-MS/MS analyses in exometabolomic experiments.

The application of NLDM to investigate the substrate preferences of 7 isolates revealed an interesting pattern of substrate utilization. Notably, all metabolites in NLDM were utilized by at least 1 isolate after 24 hours. Interestingly, sugars were among the least depleted chemical classes, with the *Cupriavidus* sp. FW305-77 not utilizing any of the sugars. In contrast, nucleosides and nucleobases were used by all isolates. Presumably, this is via purine and pyrimidine salvage pathways which are known to be major pathways for bacteria to obtain carbon, energy, and nitrogen for growth(Vogels and Van der Drift, 1976; Nygaard, 1993). Metabolite depletion patterns were similar for closely related isolates, the 4 *Pseudomonas* isolates, although differences between the individual isolates can be observed (Figure 1). The 2 most closely related isolates GW531-R1 and FW305-3-2-15-C-LB3 (with an ANI of 88%) also had the most similar utilization patterns.

The observation that all of the bacteria tested grew on NLDM suggests that this is a useful media for both exometabolite profiling and bacterial cultivation. In fact, 29 of the 53 isolates tested reached higher biomass yields on NLDM compared to R2A. All but one of the 14 *Pseudomonads* tested reached a higher biomass yield in NLDM than in R2A. This is interesting because R2A was originally developed to isolate Pseuodomonads from treated potable water(Reasoner and Geldreich, 1985). The growth rates of the isolates on R2A and NLDM were similar, indicating that the isolates have a similar fitness on both media(Ram *et al*., 2019).

A shorter lag phase was observed for some isolates on NLDM. This could be attributed to the higher concentration of immediately accessible metabolites in NLDM compared to R2A as R2A contains biopolymers which require depolymerization prior to use, resulting in delayed isolate growth(Korem Kohanim *et al*., 2018). Specifically, all sugars in NLDM are simple substrates (mono- or disaccharides) whereas half of the sugars in R2A are complex polysaccharides, which requires breakdown by extracellular glycosyl hydrolases. In addition, R2A is known to contain a diversity of peptides, which are presumably less metabolically accessible than the amino acids included in NLDM.

Compared to our previously reported soil defined medium (SDM), NLDM is richer (1146 mg/L vs 355 mg/L organic C, for NLDM and SDM respectively), has a more balanced C:N ratio (9:1 vs 1:1) and contains a wider selection of metabolites (64 vs 46 metabolites)(Jenkins *et al*., 2017). Furthermore, NLDM supported the growth of 9 isolates that did not grow on SDM(Jenkins *et al*., 2017).

There are a number of limitations to NLDM that are important to consider. As discussed above, the relative simplicity and lack of biopolymers in this media compared to soil and R2A causes its culturing characteristics to slightly differ from those of R2A. The 1x concentration of NLDM, while designed to mimic 1x R2A, is dramatically richer than soil dissolved organic carbon and nitrogen, which typically range between 0.42 to 372.1 mg C/L and 0.025 to 10 mg N/L, respectively(Ros *et al*., 2009; Langeveld *et al*., 2020).

## Conclusion

This study created a defined medium, NLDM, that enables both exometabolomic characterization and the requisite bacterial cultivation based on the metabolite composition of R2A and soil, metabolite usage by microbes, and biologically relevant elemental stoichiometry of these compounds. NLDM was found to support the growth of all of the 53 phylogenetically diverse isolates tested with comparable biomass yields and growth rates. An exometabolomics study using NLDM and 7 different isolates showed that all metabolites in NLDM were significantly depleted by at least one isolate. We anticipate that the NLDM medium will enable the examination of microbial substrate utilization for a broad range of isolates. We speculate that this media may have additional value in supporting microbial isolations and additional types of microbial characterization.

## Materials and Methods

### Chemicals

All individual chemicals were purchased from Sigma Aldrich (St. Louis, MO, USA) except for sn-glycero-3-Phosphocholine (Cayman Chemical, Ann Arbor, MI, USA). Wolfe’s Vitamin supplement (MD-VS^™^) and Wolfe’s Trace Mineral supplement (MD-TMS^™^) were purchased from ATCC (Manassas, VA, USA).

### Media preparation

NLDM was prepared by adding the 64 metabolites to a base medium composed of 1x Wolfe’s mineral and 1x Wolfe’s vitamin solutions, potassium phosphate, sodium phosphate, calcium chloride, magnesium sulfate and ammonium chloride (Supplementary Table 2). NLDM was sterilized using a 0.22um filter.

### Growth of 53 phylogenetically diverse ORFRC isolates

NLDM was compared to R2A medium at 1x concentration (Tecknova, Hollister CA) in their ability to support the growth of a broad range of ORFRC isolates, each in triplicate. For growth analysis, the 53 phylogenetically diverse isolates (Supplementary Table 6) were revived in liquid R2A medium from frozen glycerol stocks. Aliquots were washed twice with PBS and diluted with the test medium to an OD_600_ of 0.1 prior to inoculation into 96-well plates. For each fresh medium, 36 uL of starter culture was added to 144 uL test medium and plates were incubated under aerobic conditions for 24 h at 27°C, and shaken at 250 rpm. Growth data were collected by measuring OD_600_ on a TECAN Microplate reader at 15 min intervals. Growth was defined as an increase of 0.05 or greater from the first time point (max OD_600_ - initial OD_600_) after subtraction of the media control.

### Exometabolomics Sample Preparation

Quadruplicate 1 mL cultures (including medium controls) of 7 different isolates (FW305-3-2-15-E-TSA4, GW531-R1, GW456-L15, FW305-3-2-15-C-LB3, FW305-77, GW456-R20 and GW247-27LB) from the ORFRC isolate collection were cultured in NLDM at 27°C using 96 deep well plates (same inoculation technique described in the previous section). A single time point was collected from individual wells, each at 24 h. A culture fraction of 0.4 mL was centrifuged at 7,000 x g for 5 min at 4°C and 0.25 mL of the supernatant was collected. The supernatants were then frozen at −80°C, lyophilized to dryness and resuspended in 250 μL methanol. The resuspended samples were filtered through 0.2μm centrifugal filters (Pall Corporation, Port Washington, NY, USA) for 2 min at 5,000 x g and analyzed as described in the previous LC-MS/MS section.

### LC-MS/MS analysis and metabolite identification and statistical analysis

Metabolites in the NLDM were chromatographically separated using hydrophilic interaction liquid chromatography (HILIC) and detected by high resolution tandem mass spectrometry. Analyses were performed using an InfinityLab Poroshell 120 HILIC-Z column (Agilent, Santa Clare, CA, USA) on an Agilent 1290 stack connected to a Q-Exactive Hybrid Quadrupole-Orbitrap Mass Spectrometer (Thermo Fisher Scientific, Waltham, MA, USA) equipped with a Heated Electrospray Ionization (HESI-II) source probe. Separation, ionization, fragmentation and data acquisition parameters are specified in Supplementary Table 7. Briefly, metabolites were separated by gradient elution followed by MS1 and data dependent (top 2 most abundant MS1 ions not previously fragmented in last 7□s) MS2 collection; targeted data analysis was performed by comparison of sample peaks to a library of analytical standards analyzed under the same conditions. Three parameters were compared: matching m/z, retention time and fragmentation spectra using Metabolite Atlas (https://github.com/biorack/metatlas)(Bowen and Northen, 2010; Yao *et al*., 2015). Metabolite background signals detected in the extraction blanks were subtracted from the experimental sample peak heights/areas. Metabolite peak heights and peak areas were normalized by setting the maximum peak height or peak area detected in NLDM uninoculated control samples to 100%. Metabolite feature peak heights or peak areas were used for relative abundance comparisons. Hierarchical clustering analysis with a Bray–Curtis dissimilarity matrix was performed with the Python 3.7.4 Seaborn 0.9.0 package. The significance between control medium and isolate metabolic profiles was analyzed with the Python statsmodels 0.10.1 ANOVA test coupled to a Python Tukey’s honestly significant difference test with α = 0.05 corresponding to a 95% confidence level for each metabolite.

## Acknowledgements

This material by ENIGMA-Ecosystems and Networks Integrated with Genes and Molecular Assemblies (http://enigma.lbl.gov), a Science Focus Area Program at Lawrence Berkeley National Laboratory is based upon work supported by the U.S. Department of Energy, Office of Science, Office of Biological & Environmental Research under contract number DE-AC02-05CH11231.

---

**Supplementary Figure 1.**
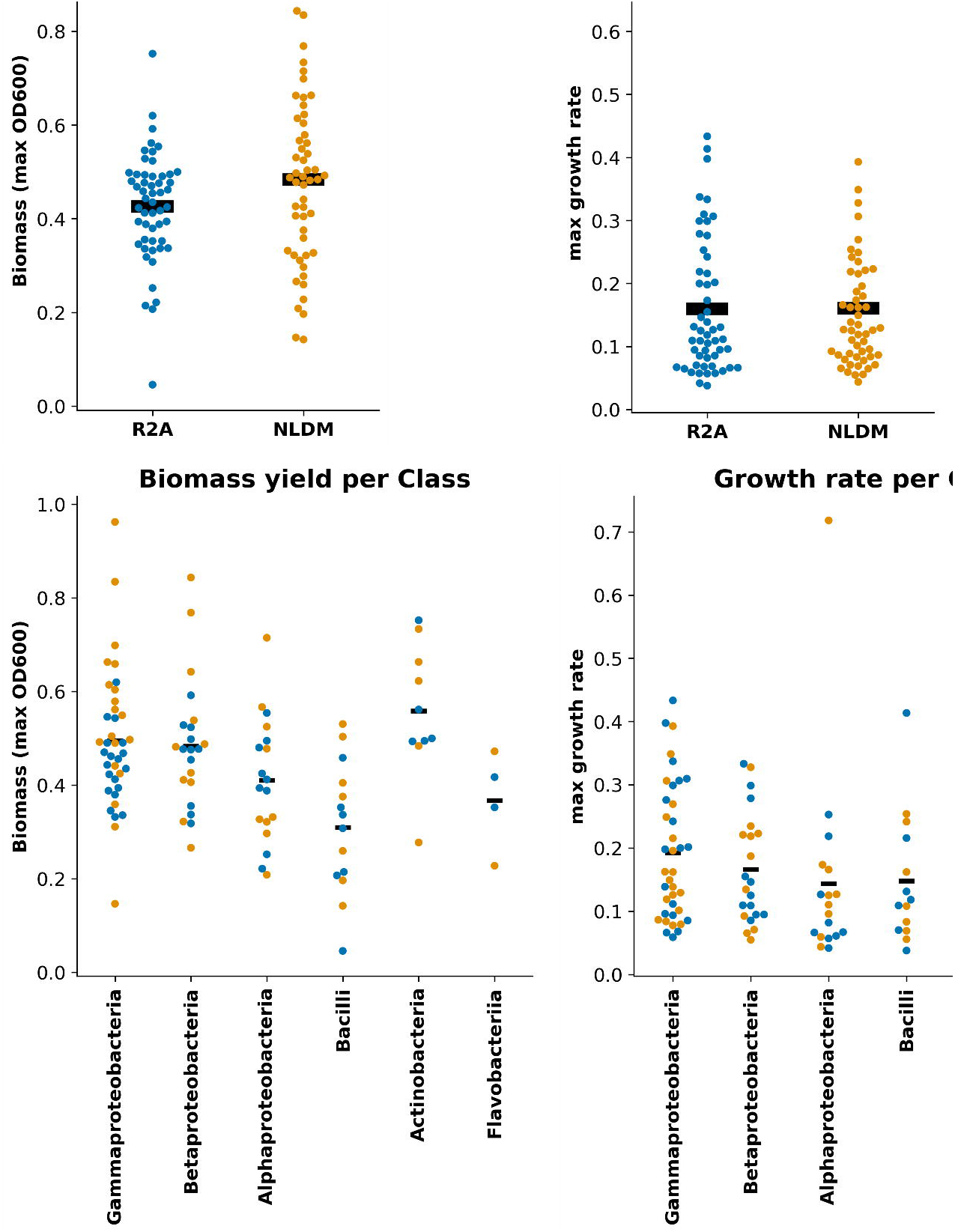
Biomass yield and growth rates of isolates grown on R2A and NLDM over 24 h.

**Supplementary Figure 2.**
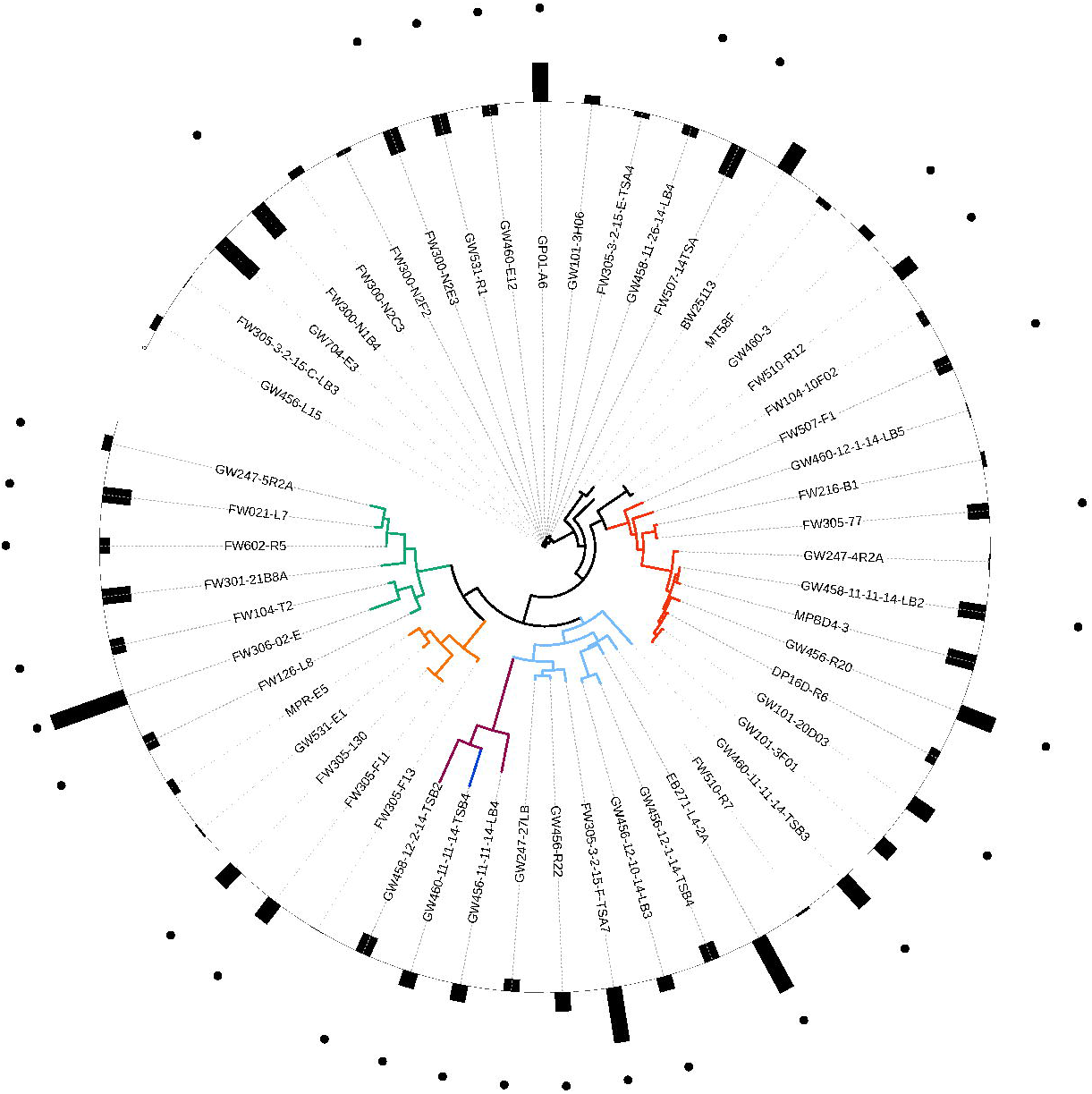
Phylogenetic tree of all isolates with corresponding ratio of growth rates on NLDM and R2A. Branch colors indicate the phylogenetic origin of each isolate by class (orange = Actinobacteria, sky blue = Alphaproteobacteria, bluish green = Bacilli, vermillion = Betaproteobacteria, reddish purple = Flavobacteria, black = Gammaproteobacteria, blue = Sphingobacteriaceae). Bars on the outer circle indicate the average (n=3) log2 ratio of the growth rate (slope of exponential growth phase) of each isolate grown on NLDM and R2A. Black dots indicate significant difference (two-sample *t*-test, *P* < 0.01). All growth data (OD_600_ values over time for all isolates) can be found in Supplementary Table 4.

